# Two Are Better Than One: Solving the Problem of Vertical Sound Source Localization via Binaural Integration of HRTFs

**DOI:** 10.1101/2020.09.10.291468

**Authors:** Timo Oess, Heiko Neumann, Marc O. Ernst

## Abstract

Early studies have shown that the localization of a sound source in the vertical plane can be accomplished with only a single ear, thus assumed the localization mechanism to be based on monaural cues. Such cues are induced by the pinna and consist of notches and peaks in the perceived spectrum which vary systematically with the elevation of sound sources. These processes pose several problems to the auditory system like identifying and extracting spectral cues on a neural level, as well as, distinguishing pinna induced peaks and notches from features already present in the source spectrum. Interestingly, at the stage of elevation estimate binaural information from both ears is already available and it seems plausible that the auditory system takes advantage of this information. Especially, since such a binaural integration can improve the localization performance dramatically as we demonstrate in the current study. For that, we first introduce a computational model architecture that takes advantage of binaural signal integration to localize sound sources in the median plane. Model performance is tested under different conditions which reveal that localization of monaural, as well as binaural inputs is best when the model is trained with binaural inputs. Furthermore, modeling results lead to the hypothesis that sound type specific prior information is taken into account to further improve localization quality. This deduced hypothesis about vertical sound source localization is confirmed in a behavioral experiment. Based on these results, we propose that elevation estimation of sound sources is facilitated by an early binaural signal integration and can incorporate sound type specific prior information for higher accuracy.

## Introduction

Audition is our only far-reaching, 360^°^ sensory system. It allows us to sense threats outside our visual field (e.g., behind us, above or occluded), thus providing a crucial factor for survival. Given that auditory signals do not contain 3D information directly, the accuracy with which humans auditorily estimate 3D locations (lateral direction, elevation, and distance) is remarkable. All such information has to be derived from the tiny vibrations of the eardrum, created by incoming sound waves of different frequency and amplitude.

Such an ability requires extensive neural computations along the auditory pathway which transform oscillations of the eardrum (tonotopic representation of the stimulus) to, e.g., comprehensible speech (phonetic representation [12, 29]) or spatial information about the location of a sound source (topographic information [4, 9]).

In order to transform tonotopic inputs of sounds to topographic information of space, the auditory system extracts specific cues from the sound signal, created by the spatial separation of the ears, their shape and the shadow of the head: The distance between the ears creates an interaural time difference (ITD) between the arrival time of the signals at the left and right ear [58]. Together with the interaural level difference (ILD), created by the attenuation of sounds by the head [45], these two cues provide a means for localization of sounds laterally.

However, for sounds with different elevation or distance that all arise from the so-called cone-of-confusion [50] these cues provide ambiguous signals. To resolve this ambiguity and to accurately localize sounds in elevation the auditory system exploits direction-dependent modulations of the proximal frequency spectrum at the eardrum relative to the distal spectrum of the sound source. Such modulations (filters) are induced by the shape of the trunk, head and ear (pinnae) [24, 36] and are reflected by troughs or peaks in the spectral filter altering the proximal sound spectrum at the eardrum. These so called spectral cues can be summarized in the Head Related Transfer Function (HRTFs, see **Fig. 1A**) [3, 5, 13, 34, 47, 55, 59]. Extracting and analyzing spectral cues from the proximal input is not straightforward as the spectrum of the distal stimulus is typically unknown and the spectra of everyday sounds are very different from each other which results in very different proximal spectra and the assumption that they share common features is not necessarily given. This poses several problems for elevation estimation:

**Figure 1.**
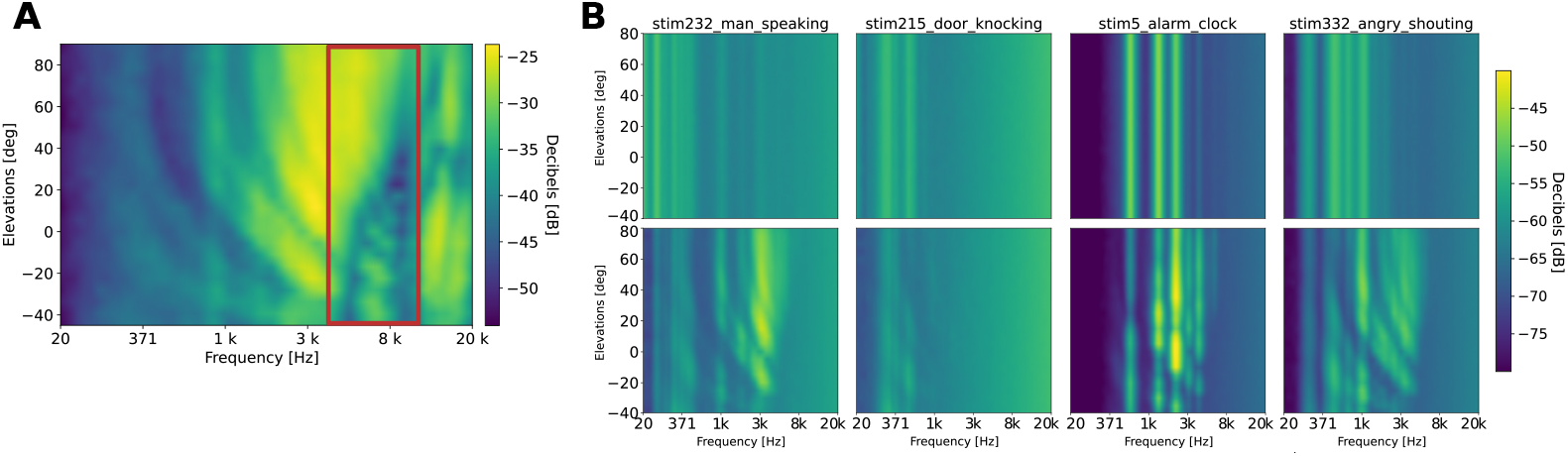
Head-Related Transfer Function. **A** HRTFs for CIPIC ^1^subject no. 10 as a function over elevations. Different colors indicate the energy content. Direction dependent changes in energy content is most prominent above 4kHz (red box). **B** Elevation spectra maps of different stimuli before being filtered with HRTF (top row). Elevation spectra maps of different natural sounds after being filtered with the HRTF of **A** (bottom row).

- At the level of the eardrum the perceived sound spectrum has already been filtered with the elevation-dependent HRTF but, in principle, the auditory system has no indication which of the spectral cues were induced by the HRTF or were already present in the source spectrum. Thus, the estimation of sound source elevation is an ill-posed problem [25, 26]. This becomes apparent when looking at the spectra of different sounds, which vary largely from each other (see **Fig. 1 B**). It is already difficult to identify the structure imposed by the HRTFs and extracting spectral cues from such highly variable input signals for learning is challenging. This necessitates a mechanism that separates the sound type specific spectral content from the HRTF induced modulations, thus leaving only the HRTF dependent frequency cues. Some computational models tried to solve this problem by taking assumptions about know certain aspects of the inputs during computation, e.g., considering a known spectrum of the sound source [38], assuming local constancy of sound spectrum [63], requiring a broadband and sufficiently flat source spectrum [33] or comparing the of left and right input signals [27].
- Another issue in vertical sound source localization is the role of binaural integration. Early localization experiments demonstrated that participants are able to localize sounds with only one ear [52] and that the ability to localize sounds with one or two ears is similar. These findings lead to the conclusion that vertical sound localization is monaural [21]. However, by using virtual sound sources provided over headphones Wightman et al. in a later study [59] questioned the monaural localization paradigm applied in most previous experiments including their own. Their findings demonstrated that localization is effectively degraded under monaural listening conditions. A later study confirmed that both ears contribute to the perception of elevation [35], thus supporting the hypothesis that binaural integration underlies the processing of elevation features in sound spectra.
- When such an integration of signals takes place is still unclear. Hofman et al. [23] first described different schemes for elevation estimation. Based on their findings they hypothesized that a weighted integration step needs to be employed for the signal from the left and right ear to derive a single estimate of elevation. Whether this integration step already takes place before the spectral-mapping or after was unclear. Later on, the authors tried to answer this question in another experiment [57] but the results are ambiguous and the authors were not able to derive a confirmatory conclusion.

The difficulty of separating HRTFs from the source spectrum, the contradicting results for monaural and binaural sound source localization [21, 59], and the unclear integration order of signals from the left and right ear [23] raise the question of how the auditory system processes sound signals on a neural level? In particular, how can it an learn a representative template of HRTFs in order to generate a stable and unique perception of source elevation estimates?

Here, we propose a model architecture of sound source localization in the median plane that extracts elevation specific cues by integrating signals from both ears and learning a sound type specific prior. The binaural integration in this architecture leads to a sound type independent representation of HRTFs, thus alleviating the ill-posed problem of elevation estimation. Based on such signals the architecture reliably localizes binaural sound sources but struggles with monaural input signals. By integrating additional sound type specific prior information for elevation estimation, the architecture becomes able to localize monaural sound signals. We first implement this architecture in an arithmetic model and conduct several experiments. Second, we provide a neural network implementation that demonstrates biological plausibility. Finally, we present a behavioral experiment in which we tested participants under monaural and binaural conditions to validate model hypotheses. Based on these results, we conclude that elevation estimation is a binaural process but can deal with monaural inputs if the sound has previously been learned, i.e., prior information is available.

## Results

In this chapter, we introduce the overall model architecture in the first section. and demonstrate its localization performance under different conditions (second and third section). The results of these sections are based on a arithmetic model implementation of the architecture (see Arithmetic model for details). In a fourth section, we introduce a neural network implementation of the architecture (see Neural model for details) and demonstrate its localization performance. The fifth and last section presents results from the behavioral experiment and confirms the model predictions.

### Brief description of the model architecture

The model architecture consists of four basic processing layers (see Fig. 2). The first layer normalizes ipsi- and contralateral inputs (I), respectively. The resulting signals from the two sides are integrated in the second layer (II). A third layer averages over all input signals thus learning a map of elevation spectra (III). In layer four (IV), perceived signals are compared to this learned map via cross-correlation (see [25] for details) to estimate the elevation of the sound source.

**Figure 2.**
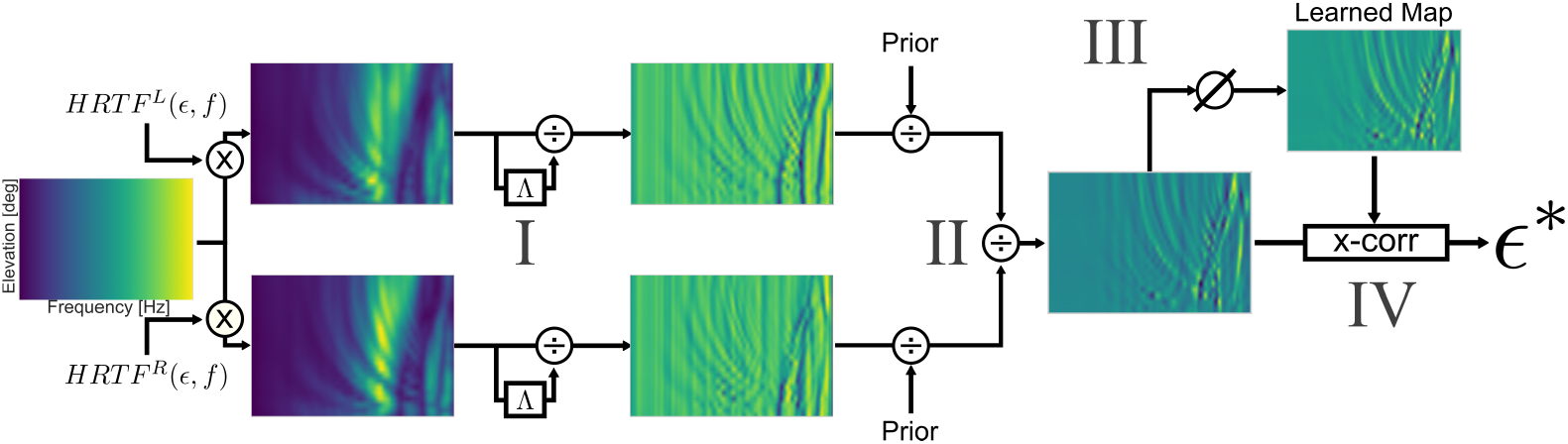
Model Architecture. Each sound that is presented to the model is first filtered by a subject’s HRTF of different elevations for the left (*HRTF*^*L*^(*E, f*)) and right ear *HRTF*^*R*^(*ϵ, f*), respectively. The resulting signals are filtered by a Gaussian normalization step (*I*). Then, if available, prior information is integrated, separately for the left and right ear. The binaural integration step (*II*) combines the signal from the left and right ear. Each perceived signal of all presented sound types contributes to build a learned map of elevation spectra (*III*) for later cross-correlation with a perceived sound to computed an elevation estimate *∈*^∗^ (IV).

The first layer of neurons receives a frequency signal of the sound signal, provided by a gammatone-filter bank [43] of the ipsi- and contralateral input signals, as an input and performs a divisive normalization with a Gaussian-filtered version of it self [8, 22]. This normalization already provides signals with prominent spectral cues. Averaging these signals over all perceived elevations leads to a sound type specific prior which is in some conditions used for map learning and to improve localization of monaural and binaural sounds. In the binaural integration layer, the normalized signals from the ipsi- and contralateral side are combined (by a division) to provide a binaural signal. In the last layer of the model, a cross-correlation of the perceived, filtered sound signal with a previously learned map is calculated and the elevation with maximum correlation value is chosen as the elevation estimate (similar to [25]). For more details on the model implementation, see Methods chapter.

We found that this model of elevation perception can account for typical human behavior [21, 30, 35] of auditory elevation perception and predicts that the underlying localization process is fundamentally binaural with the ability to localize monaural sounds when integrating prior information.

To investigate the performance and validity of our model we used HRTFs of 45 subjects from the CIPIC database [1] and presented each with 20 different natural sound types (signal-to-noise ratio 5 : 1), originating from 25 different elevations ([−45^°^, 90^°^] in 5.625^°^ steps according to CIPIC database recordings [1]). All presented sound types are averaged for each participant to create a learned map of spectral elevations. This map is used for the cross-correlation step in layer IV with a perceived probing signal which is randomly chosen from one of the previously presented sound types and elevations. The resulting elevation estimate is compared to the actual elevation of the presented sound. Consequently, for each participant a linear regression analysis is performed on this data which provides a response gain (accuracy), bias (spatial bias) and precision (coefficient of determination). Four conditions are tested to demonstrate the advantage of binaural signal integration and incorporation of prior information on localization performance:

1. **Monaural**: the binaural integration layer (*II*) is skipped and pure, normalized monaural signals are directly compared with the learned map
2. **Monaural-Prior**: again the binaural integration layer (*II*) is skipped, but the resulting monaural signal is combined with the previously learned sound type specific prior (*III*) before comparing it to the learned map.
3. **Binaural**: the binaural integration layer (*II*) remains active and resulting binaural signal is compared to the learned map for an elevation estimate.
4. **Binaural-Prior**: in this condition the binaural signals are combined with prior information (*III*) before the cross-correlation with the learned map is performed.

### A binaurally learned map can account for binaural as well as monaural sound source localization

Experimental results demonstrate that humans can localize sounds with just a single ear [21, 52]. Based on these results, the common assumption for human vertical sound source localization is that it is fundamentally monaural. That is, localization is separately initialized for the left and right ear, respectively, and the two elevation estimates are integrated for a single estimate [25]. We conduct a first experiment in which we question this assumption by demonstrating that when a combined binaural map for the left and right ear is learned, monaural sound source localization is still possible.

Here, we show that a binaurally learned map of elevation spectra can account for sound source localization under binaural and monaural conditions. However, for monaural signals, localization is only possible when prior information is integrated. Thereby, such a learned map can account for experimental results with binaural and monaural listing conditions (see section Behavioral Experiment).

Simulation results for a single participant (CIPIC HRIR no. 9) are shown in **Fig. 3 A**. When a binaural map with integrated prior information is learned, pure monaural sounds (ipsilateral ear, left panel, condition 1) are basically not localizable (gain: 0.36, bias: 28.90, score: 0.13). Surprisingly, when such monaural sounds are combined with a previously learned sound type specific prior (middle left panel, condition 2), the localization quality increases dramatically (gain: 0.62, bias: 5.63, score: 0.38), thus localization ability is restored.

**Figure 3.**
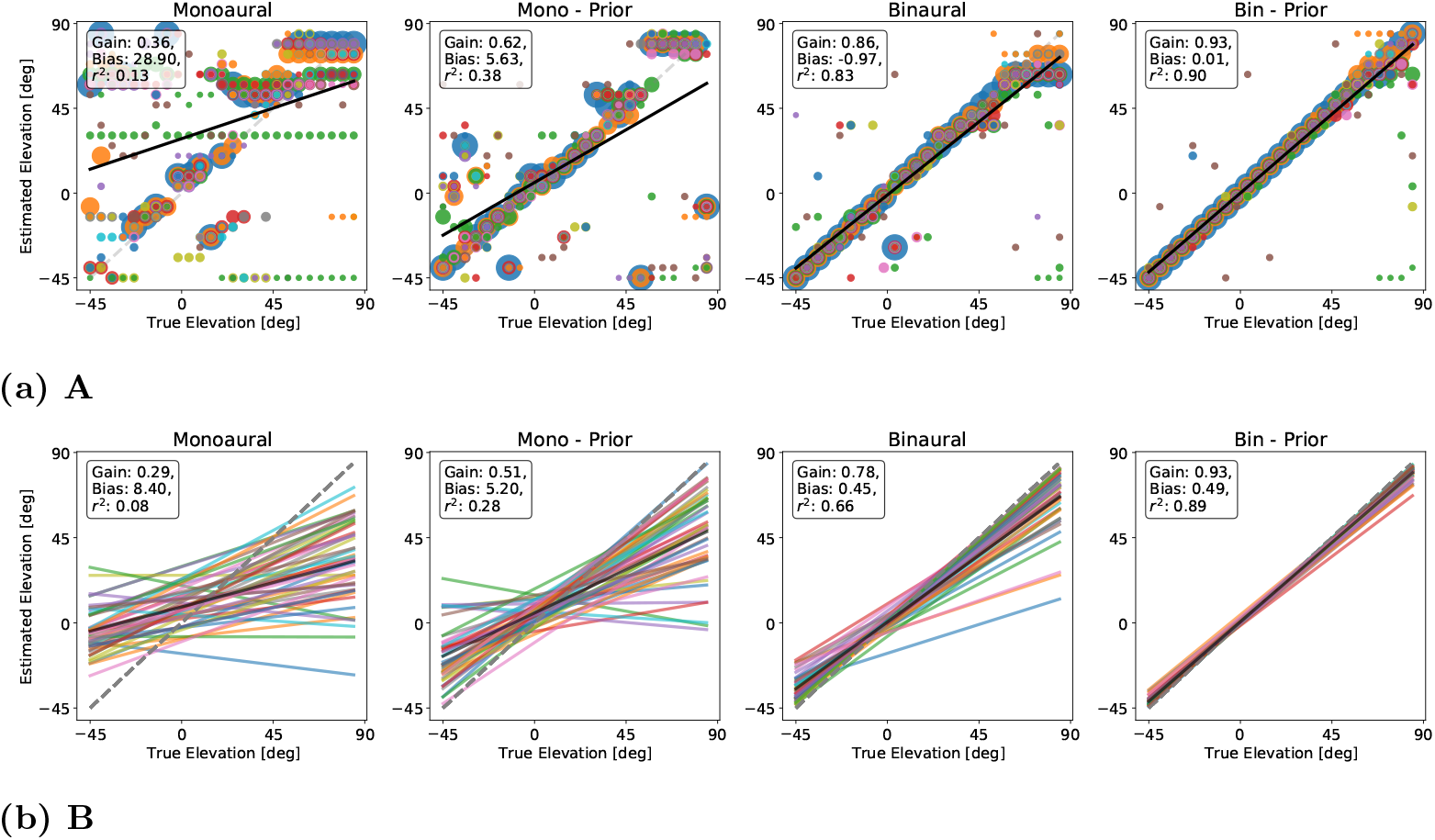
Elevation estimation results. Model estimates for all presented elevations and all sound types. X-axis indicates the elevation of the presented sound. Y-axis is the model elevation estimate for a sound. Pure monaural sounds are presented in left panel. Middle-left panel shows model estimates for monaural sounds integrating prior information. Pure binaural sounds are presented in middle-right panel. Right panel depicts model estimate for binaural sounds integrating prior information. **A** Model estimates for a single participant (no. 9 CIPIC). Each dot represents one sound source elevation with color indicating sound type. Different blob sizes are used for better visualization. **B** Calculated regression lines for different participants in the CIPIC database are shown (colored lines). Black lines are calculated by averaging over all colored lines to achieve averaged estimation values. Regression values are shown in inset box.

Localization performance for sounds that are presented binaurally with (middle right panel, condition 3) and without (right panel, condition 4) integrating prior information is almost perfect (gain: 0.86, bias: -0.97, score: 0.83 and gain: 0.94, bias: 0.01, score: 0.90, respectively). Such good performance in these conditions is expected since the learned map is constructed based on these binaural sounds integrating prior information.

When averaging localization quality over all participants (all 45 HRIRs from CIPIC database) the initial trend remains **Fig. 3 B**. That is, localization of monaural sounds is basically non-existing (left panel, condition 1, gain: 0.29, bias: 8.40, score: 0.08) but improves tremendously when prior information is integrated (middle left panel, condition 2, gain: 0.51, bias: 5.20, score: 0.28). Again, localization performance for binaural sounds and binaural sounds integrating prior information remains stable (gain: 0.78, bias: 0.45, score: 0.66 for condition 3 and gain: 0.93, bias: 0.49, score: 0.89 for condition 4 respectively).

These results demonstrate that a binaural map of elevation spectra supports the localization of monaural sounds integrating prior information but is unable to localize pure monaural sounds, since their spectral information differs greatly from the learned spectra (see supplementary Fig. S1). This is a strong indication for the existence of a binaurally learned map.

### Localization quality for differently learned maps

In the previous experiment a binaural map is learned to localize sounds signals of various types. Hofman and colleagues [23] described different possibilities of how a unique perception of signal elevation from the two ears might be achieved. They hypothesize two different schemes for elevation perception: the *spatial weighting* scheme and the *spectral weighting* scheme (see [23], their Fig. 7). The spectral weighting scheme is similar to our proposed binaural integration model with a binaurally learned map, whereas the spatial weighting scheme would correspond to our model when a monaural map is learned.

In a second experiment, we investigate which of the proposed schemes for map learning and utilization of priors is more plausible. Thereby, we validate our results of the previous experiment. Here, we test the localization quality of participants when the learned map is based on different signals (i.e. monaural, monaural-prior, binaural, binaural-prior, different rows in Fig. 4) and demonstrate that a binaural map with sound type specific prior integration produces the best localization results (Fig. 4 last row).

**Figure 4.**
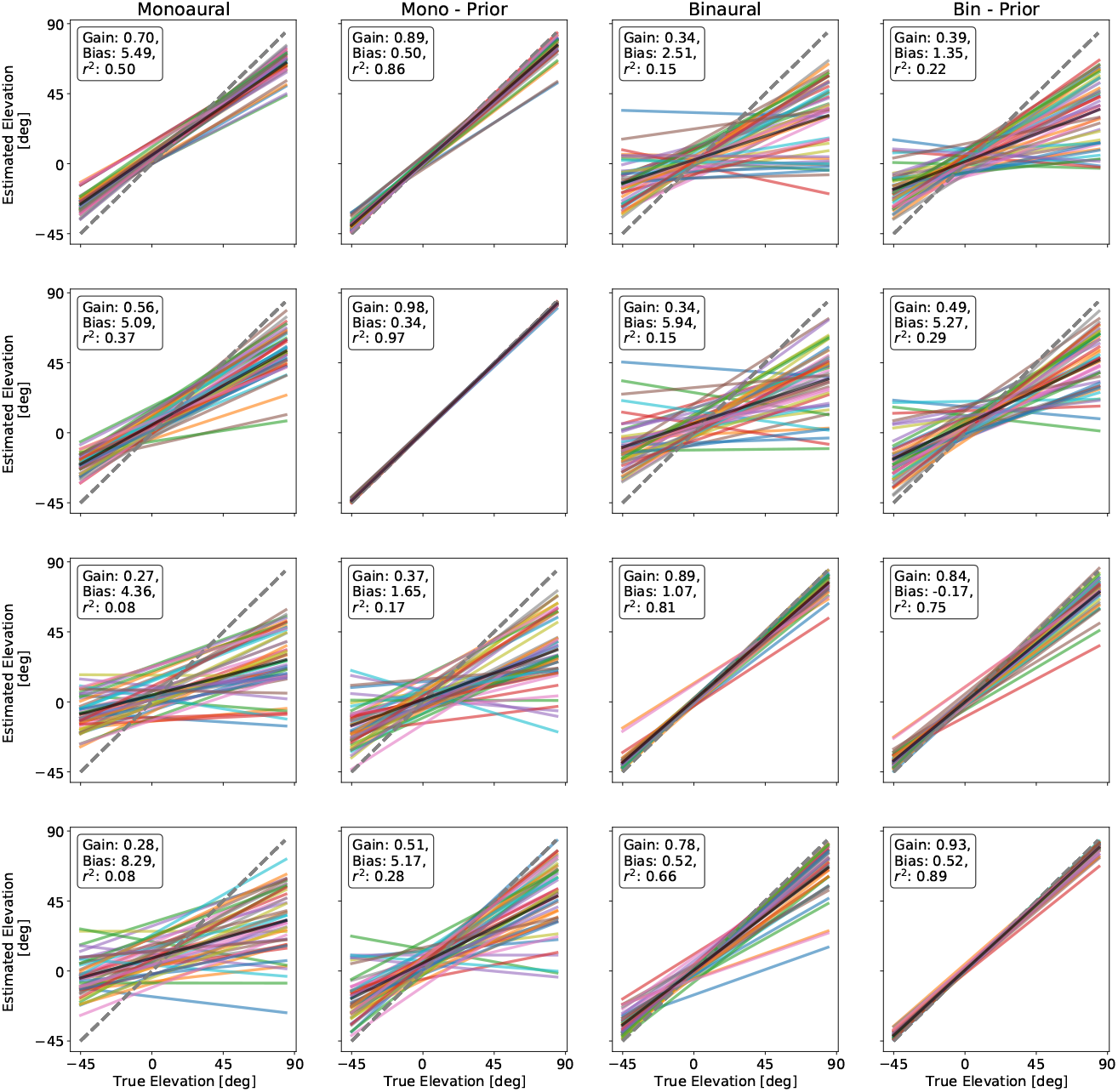
Estimation results over differently learned maps. Each column depicts localization results of the model for the different conditions, similar to Fig. 3. X-axis indicates the elevation of the presented sound. Y-axis is the model elevation estimate for a sound. Pure monaural sounds are presented in left panel. Middle-left panel shows model estimates for monaural sounds integrating prior information. Pure binaural sounds are presented in middle-right panel. Right panel depicts model estimate for binaural sounds integrating prior information. In each row the learned map, which is used to localize a sound, is learned based on different signals. In the first row, the map is based on pure monaural sounds. Monaural sounds integrating prior information are used to build the learned map for the second row. In the third row, the map is based on pure binaural sounds. Binaural sounds integrating prior information are used to build the learned map for the fourth row.

Our investigations show that when the learned map of elevation spectra is based of pure monaural input signals, localization performance is best for monaural signals integrated with sound type specific prior information (gain: 0.89, bias: 0.50, score: 0.86, condition 2). These sound signals are even better localized than pure monaural signals (the basis for the map, gain: 0.70, bias: 5.49, score: 0.50, condition 1). This demonstrates the benefit of the integration of sound type specific prior information. This advantage of prior integration can be also seen in the binaural sound conditions. For pure binaural sounds the localization performance is worse compared to binaural signals integrating prior information (gain: 0.34, bias: 2.51, score: 0.15 for condition 3, and gain: 0.39, bias: 1.35, score: 0.22 for condition 4, respectively). Here, the binaural prior condition even outperforms the pure monaural condition, which is surprising since binaural signals differ substantially from monaural signals (see supplementary Fig. S1).

Taken together, these simulation results demonstrate that a pure monaural map is not sufficient to localize pure monaural sounds (Fig. 4, first row, first column). Prior information is required to localize sounds in monaural and binaural conditions. Even if this prior information is integrated in the learned map (Fig. 4, second row), localization of pure monaural sounds is difficult and again prior information of the input signals is crucial. However, if the learned map is based on binaural signals with or without the integration of prior information (Fig. 4, third and fourth row, respectively) localization performance for each condition except the pure monaural condition is close to optimal. Thus, we hypothesize that elevation estimation is essentially based on binaural signals but can deal with monaural signals when prior information of such signals is available. Furthermore, these results demonstrate that prior information of sounds consistently improves localization performance of sound sources.

### Neural network model

To demonstrate the biological plausibility of the architecture, we developed a neural network model which incorporates key functionalities specified in the architecture and investigate its performance on localizing sound sources, similar to experiment one. In this implementation, a single compartment model neuron with gradual activation is employed. The activations represent the average potential of a group of real neurons in a column of brain tissue. We define such activities according to a leaky integrator equation that models membrane potential dynamics with the membrane current as the sum of excitatory, inhibitory, and leak conductances [11, 16, 28] (see Methods for details). The neuron populations are implemented and connected with each other according to the different layers defined in the arithmetic model. In the following experiment the signal-to-noise ratio is set to 0.

For this third experiment, the localization performance for all simulated participants from the CIPIC database is presented in Fig. 5. Even though different in the linear regression values the overall trend of the localization quality in the different conditions is similar to the arithmetic model. Pure monaural sounds (ispilateral ear, left panel, condition 1) are basically not localizable (gain: 0.28, bias: 2.89, score: 0.09). When combined with prior information such monaural sounds (middle left panel, condition 2) the localization quality increases (gain: 0.40, bias: 1.29, score: 0.19). For binaurally presented sounds (condition 3) the localization performance is again improved (gain: 0.67, bias: −1.82, score: 0.51) and for binaural sounds integrating prior information (condition 4) the localization performance is close to the arithmetic model (gain: 0.87, bias: −0.76, score: 0.81).

**Figure 5.**
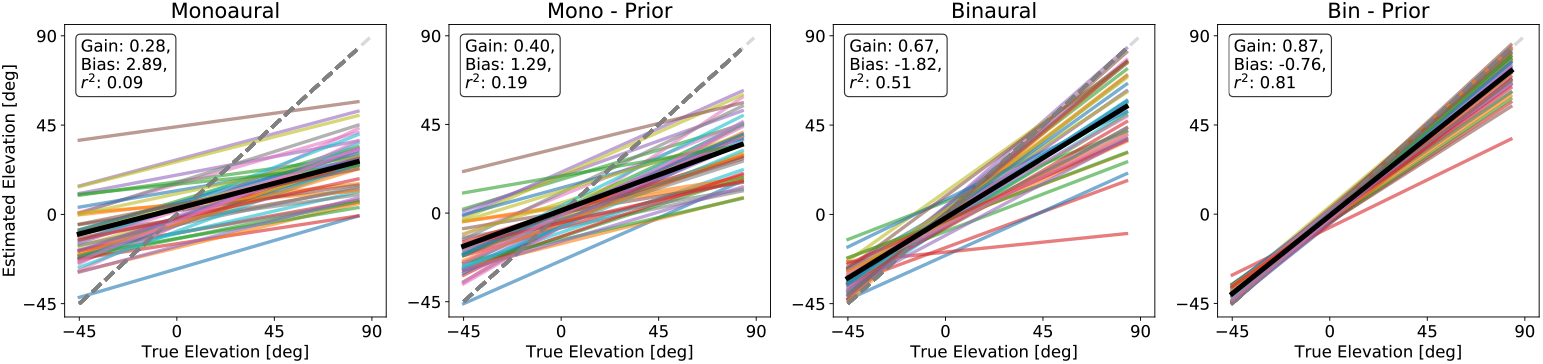
Neural Network Elevation Estimates. Model estimates for a single participant (no. 8 CIPIC). Calculated regression lines for different participants in the CIPIC database are shown (colored lines). Black lines are calculated by averaging over all colored lines to achieve averaged estimation values. X-axis indicates the elevation of the presented sound. Y-axis is the model elevation estimate for a sound. Pure monaural sounds are presented in left panel. Middle-left panel shows model estimates for monaural sounds integrating prior information. Pure binaural sounds are presented in middle-right panel. Right panel depicts model estimate for binaural sounds integrating prior information. Regression values are shown in inset box.

### Behavioral Experiment

To confirm our model predictions, resulting from the previous simulation experiments, we conduct a behavioral experiment in which human participants (n=8) are asked to localize sound sources on the median plane. The experimental setup consisted of 11 speakers aligned on a half circle with the participant’s head in the center to ensure equal distance (1.20*m*??) from the head to each speakers (see Supplementary Fig. XX). We replicated the four different conditions of our model experiments by presenting two different sound types (white noise and rippled noise), that are assumed to be previously known (white noise) and unknown (rippled noise), respectively (for creation of sounds, see Methods). Each of these sounds is presented under monaural and binaural listening conditions. That is, participants wear an earplug and noise-canceling headphones over the non-dominant ear to prevent the perception of any sound in the monaural listening condition, whereas both ears are free in the binaural condition. Thereby, 4 distinct listening conditions, similar to the model experiments, are created:

1. **Monaural**: Rippled noise (no prior information available) is presented with the non-dominant ear occluded.
2. **Monaural-Prior**: White (prior information available) noise is presented with the non-dominant ear occluded.
3. **Binaural**: Rippled noise (no prior information available) is presented with both ears free.
4. **Binaural-Prior**: White (prior information available) is presented with both ears free.

For more details on experimental procedure and data analysis see Methods.

For this fourth experiment, the localization performance for all participants is presented in Fig. 6. Localization performance of listeners in the pure monaural condition (left panel) is low (gain: 0.25, bias: 22.56, score: 0.27), which confirms the prediction of the model that when sounds are localized with just a single ear without having access to prior information of that sound, localization is not possible. However, when prior information is available (condition 2, middle left panel), localization performance increases (gain: 0.50, bias: 18.02, score: 0.47). For binaural conditions 3 & 4 (middle right and right panel) without and with prior information available localization performance is restored (gain: 0.74, bias: 23.58, score: 0.80 and gain: 0.81, bias: 19.86, score: 0.86, respectively). As in model simulations, localization performance consistently increases when prior information is available.

**Figure 6.**
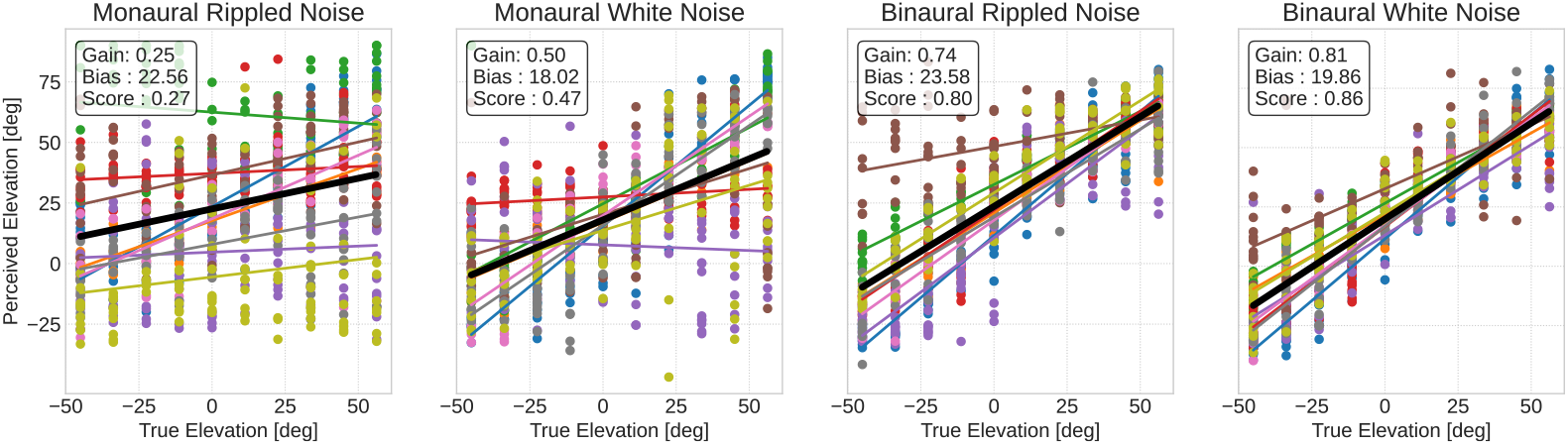
Behavioral Results. Black lines are calculated by averaging over all colored lines to achieve averaged estimation values. X-axis indicates the elevation of the presented sound. Y-axis is the participants’ elevation estimate for a sound. Elevation estimates for the monaural condition (one ear occluded) for rippled noise (no prior) in left panel. Middle-left panel shows model estimate for the monaural condition for white noise sounds (prior available). Results for binaural condition (both ears free) for rippled noise (no prior) are shown in middle-right panel. Right panel depicts model estimate for binaural condition (both ears free) for white noise (prior available). Regression values are shown in inset box.

## Discussion

In this study we presented a possible solution to the ill-posed problem of sound source localization. That is, we demonstrated that binaural signal integration leads to a vast improvement of localization performance in both modeling experiments as well as behavioral results. The proposed architecture for sound source localization in the vertical plane allows the localization of monaurally as well as binaurally presented sounds based on a binaurally learned map. In addition, we proposed sound type specific prior information which constantly improves the localization and accounts for early research that demonstrate that a monaural signal is sufficient for localization. If only monaural signals are available sound source localization remains difficult. However, when monaural sound type specific prior information is integrated, localization performance of monaural sounds is restored.

### Implications on the current view of vertical sound source localization

Our findings are in contrast to the results of [21], in which the authors presented participants with white noise and presumably unfamiliar filtered noise to test the localization performance under monaural and binaural conditions for known and unknown sounds, respectively. The authors’ results indicate that the additional binaural information in the binaural condition does not improve localization performance and that known and unknown sounds are localized equally well. Our results, on the other hand, clearly indicate that unknown sounds are basically impossible to localize under monaural conditions. Thus, we believe that the behavioral results of [21] are misleading because of two major factors: The method to occlude one ear might not be sufficient to ensure pure monaural information, as already pointed out in [60]. The findings of Wightman and colleagues [60] questioned the results of several previous experiments on monaural localization performance and demonstrated that when participants are presented with a pure monaural signal over headphones, localization is basically not possible. The authors suggested that in previous experiments with contradicting results, the occlusion of one ear was not sufficient to block all informative signals or that small head movements have been used to localize a sound. This is inline with our results from the first experiment which demonstrates that localization is essentially eliminated for monaural sounds without integrating any further information. The second factor is the choice of the unknown sound, which is a filtered white noise stimulus with random peaks and notches similar to the ones provided by the HRTF. However, such a white noise stimulus is not necessarily an unknown sound but might merely lead to a confusion between sounds from different elevations.

In order to manifest our prediction, we implemented a behavioral experiment in which we tested participants with known and unknown sounds under monaural and binaural conditions. Our model architecture predicts that unknown monaural sounds will be very difficult to localize. Though, unknown binaural sounds should be localized accurately and quickly. The behavioral results presented confirm these model predictions.

Hofman and van Opstal [23] already suggested that the elevation estimation is facilitated by a binaural interaction of the left and right ear. They introduced two conceptual schemes for this interaction, the *spatial* and *spectral* weighting scheme (see second experiment). However, it is still unclear which of these schemes is applied [57]. Our model architecture and results from the second experiment demonstrate that binaural integration is most likely taking place before the computation of an elevation estimate (spatial mapping stage), since it enables the system to extract unique elevation dependent cues and remove unnecessary source spectrum induced spectral information.

The process of binaural signal integration is an integral part of *horizontal* sound source localization and provides cues like interaural level or time difference. The fundamental principles used for the computation of these two cues are similar and are based on the integration of excitatory inputs from the ipsilateral side and inhibitory input signals from the contralateral side [6, 18, 19, 61]. It is therefore plausible that the process of binaural integration, as shown in our model, is adopted to provide distinct cues for vertical sound sources.

### Prior Information

Another major finding of our model is that the integration of sound type specific prior information facilitates monaural sound source localization as well as it improves binaural localization performance. By learning sound type specific prior information, which consists of the mean frequency components over elevations, localization performances for all conditions are improved (see Fig. 4). In our model, we assume that this prior information is learned in higher stages of auditory processing, which are able to identify a sound or at least categorize it [2, 20]. Consequently, these stages presumably provide such prior information by feedback connections to lower sensory processing areas like the inferior colliculus [32, 44, 49]. If this is the case one could measure a difference in localization speed between monaural and binaural sounds, since monaural sounds can be localized only after they have been categorized in a higher stage and a feedback signal has been sent back to the sensory stage. Nevertheless, binaural signals can be localized immediately without the use of prior information, the prior information just increases accuracy.

### Neural implementation

In a last experiment we introduced a neural implementation of the presented architecture, that implements different neuron populations and interactions of excitatory and inhibitory signals among them to replicate computations of the arithmetic model in a biologically plausible fashion. In [37] the authors investigated typical responses of neurons in the dorsal cochlear nucleus to stimuli with spectral notches and discovered that these neurons already show a sensitivity to spectral notches. Our proposed model is similar to their type II and type IV neurons in a sense that it receives excitatory inputs from the best frequency of a neuron and inhibitory inputs from neighbouring frequency bands (wide-band inhibition, see Fig. 7). Similar investigations of the inferior colliculus have found neurons that specialized in processing directional dependent features of the HRTF [10]. Our neural model follows these findings and additionally, incorporates inhibitory input to neurons in the inferior colliculus from neurons at the contralateral side to enable binaural integration. Such inhibitory connections between contralaterally placed neurons have been shown to exist [49]. The fact that in our neural model identical neuron parameters are used for all participants demonstrates on the one hand the robustness of the model. On the other hand it provides an option to improve the performance by tuning the neuron parameters specifically for each participant.

**Figure 7.**
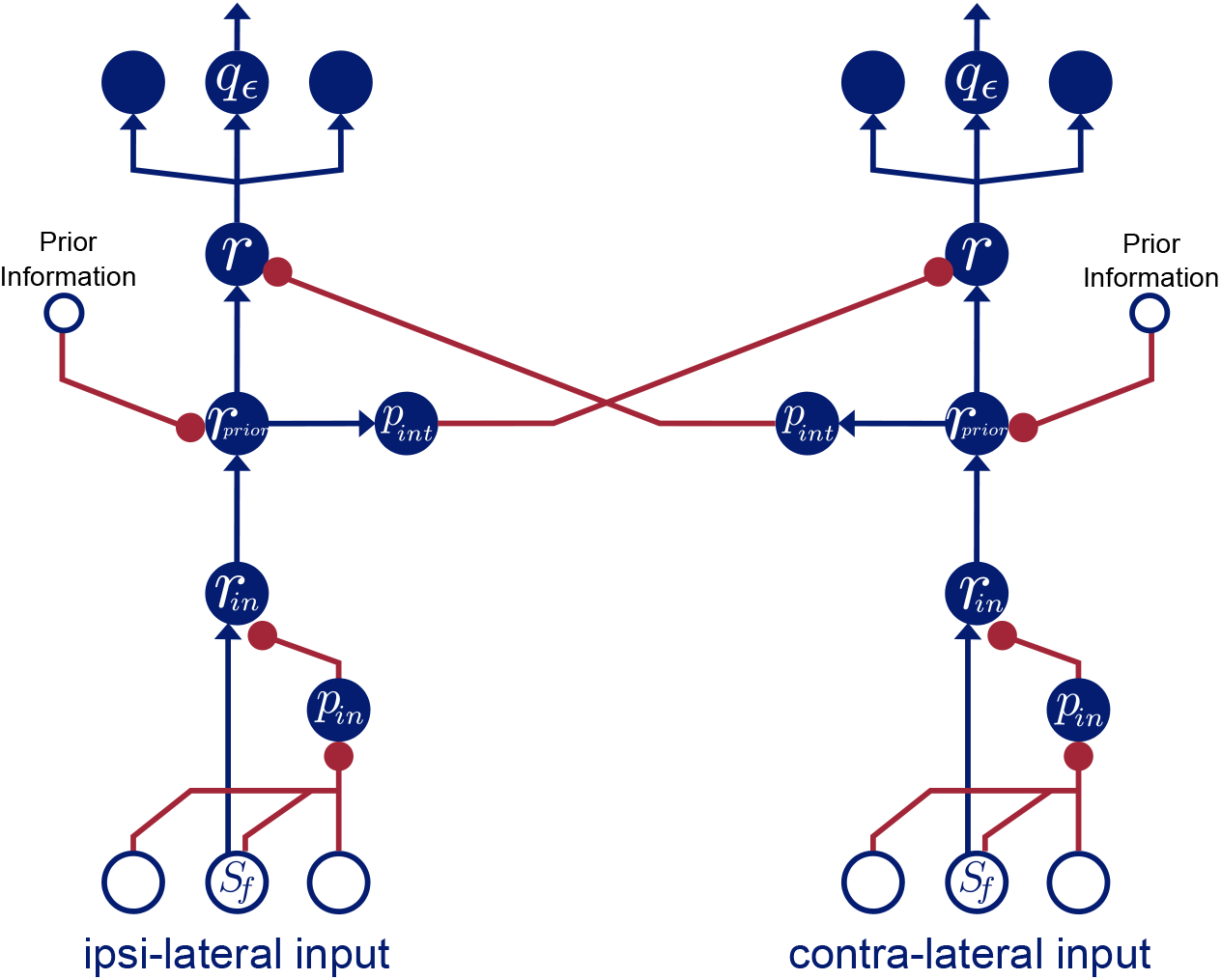
Neural Network Architecture. Blue filled circles indicate model neurons. Blue empty circles represent model inputs. Blue arrow-headed connections are excitatory and red bullet-headed connections are inhibitory connections from inputs to neurons and from neurons to other neurons, respectively. The processing consists of two independent parallel pathways for the left and right ear input which interact at the level of the superior olive and presumably inferior colliculus.

In addition to these experimental results, the structure of the model also provides a hint on when the integration of the signals from the left and right ear are integrated. Therefore, the proposed architecture for binaural integration offers an excellent basis for understanding vertical sound source localization and guides future behavioral and physiological experiments.

The presented experiments for monaural and binaural sound source localization challenges the current view on the fundamentals of how listeners localize the vertical component of environmental sounds. We propose that vertical sound source localization takes advantage of binaural signal integration in every day situations but is also capable of localizing monaural signals providing they have been heard (learned) beforehand.

## Methods

### Input data creation

Inputs 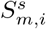 to the model are generated by, first, convolving a mono sound signal *x*_*i*_(*t*) of sound type *i* with recorded head-related impulse responses (HRIR), separately for the ipsi- and contralateral ear, denoted by *s* = *ipsi* and *s* = *contra*, respectively, of listener *m* provided by the CIPIC database [1] to model simulated sound signals arriving at the cochlea

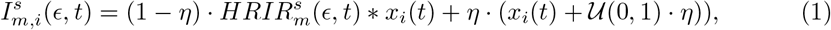

where * is the convolution of two signals, *U* (0, 1) the uniform distribution and *η* describes the signal-to-noise ratio and is commonly set to 0.2. The input noise of the data is modeled so that a part of the original, unfiltered signal (*x*_*i*_(*t*) in second term) is perceived together with random noise (𝒰 (0, 1)). For the influence of the signal-to noise ratio parameter on the localization ability see supplementary Fig. 2.

The cochlea response over frequencies for a perceived sound signal can be simulated using gammatone-filter banks [43]. This transformation from time into frequency domain is implemented by using a python implementation (https://github.com/detly/gammatone) of the auditory toolbox [51] with 24 frequency bands (replicating critical bands in human hearing [64]), window length of *twin* = 0.1*s* and step time 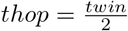. Thus, each signal 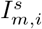 at the eardrum is transformed to its frequency domain representation by

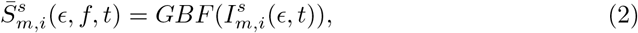

where *GBF* is the gammatone-filter bank as described in [51]. The resulting spectrum is set to be in range [100, 20000]*Hz*.

Note that this is rather an approximation of the rate-code forwarded by the auditory nerve. This approximation does not take into account rate saturation, two-tone suppression or other modifications of the spectrum by the cochlea. Nevertheless, this approach has been demonstrated to be sufficient since it preserves the differences in the spectrum of stimuli from different locations (see [38, 62]).

After this filtering step, the log power of the signal is calculated by 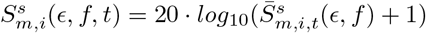. This power spectrogram is averaged over time for the final spectrum of the perceived sound

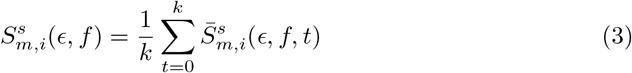

with *k* the number of time steps calculated by the gammatone filter bank.

To provide signals with similar energy levels each spectrum is normalized over frequencies:

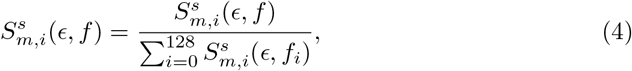

These transformation steps are separately initiated for each listener (45 HRIR from the CIPIC database), each sound type (in total 20 different sounds) and each elevation (25 in total) ranging from [−45^°^, +90^°^] in 5.625^°^ steps on the median plane.

### Sounds

Sounds that are used for the presented experiments can be found under http://mcdermottlab.mit.edu/svnh/Natural-Sound/Stimuli.html and have been previously used in other studies (see [39]. We randomly chose 25 stimuli from this set to be tested with our architecture.

## Model Description

Two different version of the binaural integration model were simulated: a arithmetic model that uses a sequence of mathematical operations and a neural model that is based on different neuron populations implementing similar operations as the first model. Response of each neuron in such populations is described by a first-order differential equation of its membrane potential. This model is provided to demonstrate the biologically plausibility of our model. If not stated otherwise all presented results are based on the arithmetic model.

### Arithmetic model

The basic model consists of three consecutive processing stages which are hierarchically organized into layers with an optional layer for the integration of prior information. This additional layer is used only in “prior” conditions or when a prior integrating map is learned.

The first layer in the model is a normalization layer that receives the frequency signal 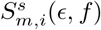 as an input and normalizes it with a Gaussian-filtered version of itself

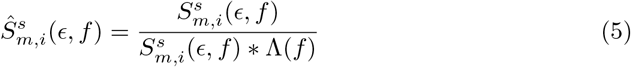

where Λ(*f*) is a Gaussian kernel with *σ* = 1. This normalization already provides signals with prominent spectral cues.

The optional prior integration step calculates a sound type specific prior by averaging these filtered signals over all elevations (n=25):

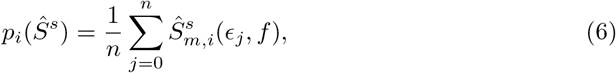

Such prior information is used in the “prior” conditions to effectively remove sound type specific peculiarities in the perceived frequency spectrum. Thus, enabling monaural localization. This additional source of averaged prior information is combined with the filtered sound signal by a simple division 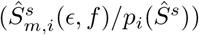. Note, that this step is omitted for conditions in which no prior information is considered.

These signals from the ipsi- and contralateral side are combined (by a division) in the integration layer to provide a binaural signal *S*^*b*^(*∈, f*):

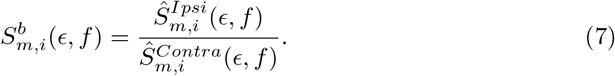

This step effectively removes sound type specific information in the signal so that only HRTF induced frequency modulations remain, making it simple for the model to localize such signals. The resulting signal is normalized over frequencies to ensure values in a feasible range 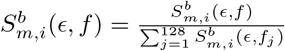.

Finally, the output layer performs a cross correlation of either 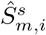 for the monaural (omitting the prior integration step) and monaural-prior conditions or 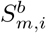 for the binaural (omitting the prior integration step) and binaural-prior conditions with a previously learned map *M*_*m*_(∈, *f*) to estimate the elevation ∈^*^ of the perceived sound source

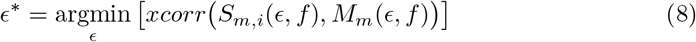

The learned map *M*_*m*_(*∈, f*) for a participant is previously constructed by averaging over all presented sound types

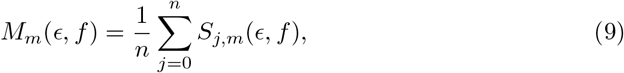

here, *S* again depends on which condition is tested. For the monaural and monaural-prior conditions *S* = Ŝ ^*s*^ and for the binaural and binaural-prio conditions *S* = *S*^*b*^.

### Neural model

The following neural model for elevation estimation is based on the computational layers introduced with the arithmetic model (see Fig.7). Each layer is realized with one or two populations of *N* neurons selective to frequency band *f* which are modeled by a first-order differential equation of the neuron’s membrane potential. This membrane potential is transformed to a firing rate by an activation function *g*(•) which is a simple linear rectified function with saturation

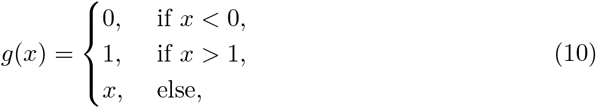

with saturation level of 1.

The core of the model is an integration population that receives excitatory input from neurons of the ipsilateral side and inhibitory inputs from neurons of the contralateral side, thus performs a binaural signal integration

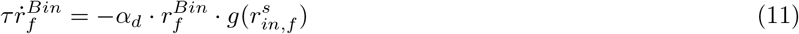

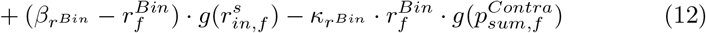

Here, parameter *τ* defines the membrane time constant, *α*_*d*_ is a default passive membrane leak conductance, 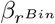 describes a saturation level of excitatory inputs and 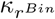 define the strength of divisive influence of the inhibitory input. A special feature of the neuron is that the the input modulates the decay rate of the neuron so that higher inputs lead to a faster decay which leads an alignment of signals with very different intensities. Such a generic neuron model has previously demonstrated to resemble typical neuronal response and to successfully solve a variety of tasks [31, 40, 41, 53]. Since such a model approach does not lead to specific voltage traces of neurons the nomenclature differs from typical electrophysiological descriptions but is in line with previous computational models [7, 17, 42, 48].

The inhibitory input 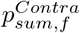 is modeled by an intermediate inhibitory population of neurons of the contralateral side

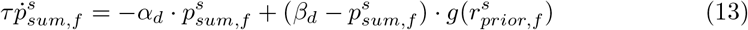

The input 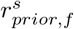 to such neurons is provided by a population of so called prior integration neurons and is modeled by

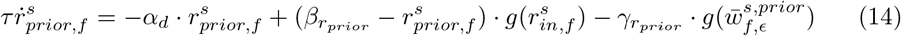

These neurons receive, presumably, cortical inhibitory input which is the mean over elevations based on a previously learned, sound type specific signal 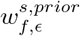 for the ipsi- and contralateral side, respectively.

Similarly, the excitatory input 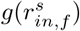 is modeled by neurons at side *s*

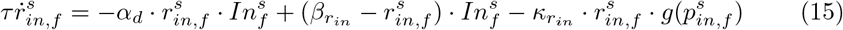

This population of neurons in the neural model realizes the Gaussian normalization layer of the arithmetic model by integrating inhibitory inputs from an inhibitory input population

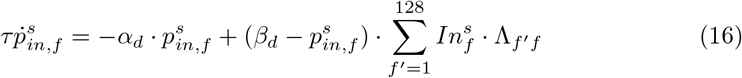

The input kernel Λ_*f ′ f*_ enables an integration of inputs over several frequency bands *f* and is defined as 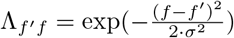 with *σ* = 3.

To ensure valid input values to the neural model, the input 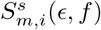 over frequency band *f* for a single participant *m*, elevation *∈* and a sound type *i* is normalized by

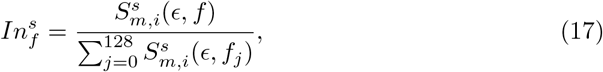

again *s ∈ {Ipsi, Contra}* depending on the input side.

To estimate the elevation of a perceived sound source the final readout layer of the network is defined as a set of 25 neurons 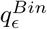, each tuned to a certain elevation ∈

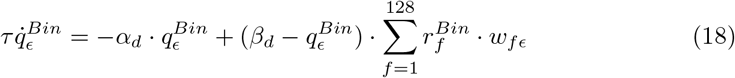

It receives excitatory inputs from the binaural integration layer and integrates them according to a previously learned weight kernel *w*_*fc*_. For an elevation estimate the index ∈* of the neuron with maximal activity is determined

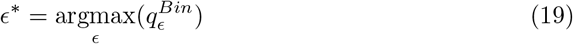

All presented results of the neural network model are calculated from the network responses readout at a single neuron level after keeping the input stimuli constant for at least 3000 time steps. This duration is sufficient for the neuron to dynamically converge to its equilibrium membrane potential of numerical integration of the state equations. For the numerical integration of the state equations we chose Euler’s method with a step size of Δ*t* = 0.0001 (for details see [54]).

The weight kernel *w*_*f ϵ*_ is learned by a supervised learning approach, where the connection weights are strengthened only when they are different from the input signal (pre-synaptic), similar to instar learning [15]. However, in contrast to pure instar learning, the learning process applied here is supervised in a sense, that adaptation of connection weights only takes place for an active visual guidance signal *v*

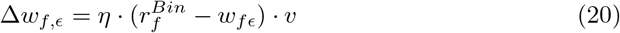

where *η* = 0.00005 is the learning rate and *v* is a vector of 25 entries, one for each elevation and is assumed to provide a visual guidance signal. That is, for a sound signal arriving from elevation ∈ entry *v*_*∈*_ of the vector is set to 1 while all other entries remain 0.

The sound type specific prior is learned separately for the ipsi- and contralateral side and is based on the activation of the prior integration neurons:

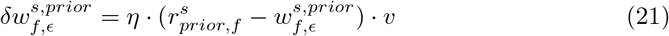

here, the values of *η* and *v* are set as described above.

The learning phase consists of 15000 trials. In each trial a sound signal from a randomly chosen elevations and sound type is presented to the model. After this learning phase the weights are normalized over frequencies to ensure similar energy content 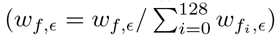 [14, 56]. Subsequently, localization performance is tested by presenting all sound signals to the model and calculating the elevation response ∈^*^. For this localization phase *η* is set to 0 to disable learning.

**Table 1.** Model parameters

| General Parameters |  |  |  |
| --- | --- | --- | --- |
| N (# Neurons) | 128 |  |  |
| $\sigma_{Kernel}$ | 3 | | |
| $\tau_d$ | 0.005 | $\alpha_d$ | 1.0 |
| $\beta_d$ | 1.0 | | |
| Excitatory Input Neuron $r_{in}$ | | | |
| $\beta_{r_{in}}$ | 200.0 | $\kappa_{r_{in}}$ | 200.0 |
| Prior Integration Neuron $r_{prior}$ | | | |
| $\beta_{r_{prior}}$ | 2.0 | $\gamma_{r_{prior}}$ | 1.0 |
| Integration Neuron $r^{Bin}$ | | | |
| $\beta_{r^{Bin}}$ | 2.0 | $\kappa_{r^{Bin}}$ | 1.0 |

## Behavioral Experiment

The results of the presented model architecture predicts a binaural integration of input signals for better localization of sound sources. In addition, we demonstrated that for monaural sounds, localization is still possible if sound type specific prior information is available. To test these model predictions, we designed a behavioral experiment in which we probe the localization performance of human participants under monaural and binaural listing conditions.

### Setup

The experimental setup is located in a sound attenuated room and consists of 13 loudspeakers (Genelec 820c, linear frequency response), arranged on an arc around the participant (see Supplementary Fig. S3 **A**) to ensure a constant distance of 120*cm* from the center of the participant’s head. The highest speaker in terms of elevation is located at 90^°^ on the median plane, i.e., vertically above the participant. From there on, each speaker is spaced 11.25^°^ apart from the previous one. The speakers are connected to a digital amplifier (RME Fireface 802) which allows for separate control of each speaker. The output volume of each speaker is equal and validated with a sound level meter before the actual experiment, placed at the position of the participant’s head.

A participant is place in the center of the speaker arc with an equal distance to all speakers. The head of the participant is placed on a chin rest to keep it fixed during the entire experimental session. The participant is asked to use a pointing device to point at the perceived location of a sound. A red dot indicates where the participant is pointing at. Despite of this red dot, the room is completely dark to prevent any visual bias. By pressing a button on the pointing the device the response (estimated location of the sound source) is confirmed and recorded.

### Conditions

Before the start of the experiment, the dominant ear of the participant is determined [46]. After that, the participant has to wear earplugs and headphone on both ears. The perception threshold is determined by playing sounds of different volume from the speaker at 0^°^ and asking the participant whether they could perceive a sound or not. The resulting volume is used throughout the entire experiment.

To align with the tested conditions of the presented modeling results, participants are asked to localize two distinct sounds (rippled noise and white noise) under monaural and binaural listening conditions. Each of these two conditions consists of 200 trials. In the monaural condition, the participants are wearing earplugs and headphones on the side contralateral to the dominant ear. In the binaural condition both ears are free. The starting condition of the experiment is chosen randomly. We tested 4 participants with monaural starting condition and 4 participants with binaural starting condition (total number of participants n=8). The participants have a 5*min* break between the conditions. In each trial one of the speaker location is chosen randomly and a randomly chosen sound is played back from this speaker. Once a sound is played back, the task of the participant is to point to the direction from which the sound is perceived as fast as possible and press the confirmation button. After each button press (trial), there is a pause of 1*s*, which is followed by another sound. We ensured that each speaker is chosen 10 times and that the distribution of chosen sounds is equal, i.e., each sound is chosen 100 times in each condition.

The two presented sounds have a duration of 300*ms* with 5*ms* linear onset and offset ramps. The white noise sound has a flat spectrum. In contrast, for the rippled noise sound a flat spectrum serves as a basis but is interleaved with artificially generated drops of energy level for specific frequency band windows (width 1000*Hz*). That is, the energy for a frequency region within the window is reduced by 95%, then the window is shifted by 2000*Hz* to the next region. This is repeated until the window reaches 20.000Hz (see Supplementary Fig. S3 **B**). By abruptly (sharp edge) reducing the energy for some frequency bands, we created a deeply artificial sound for which we assume participants do not have any prior information available. Both sounds are generated by a python script.

In the presented experiment, only the lower 10 speakers are used corresponding to the extension of the visual field of the participant. This results in the following locations of the speakers on the arc : +56.25^°^, +45^°^, +33.75^°^, +22.50^°^, +11.25^°^, 0^°^, −11.25^°^, −22.50^°^, −33.75^°^, −45^°^. All experiments have been approved by an ethics committee (approval no.104/20).

## Footnotes

1 “CIPIC HRTF Database is a public-domain database of high-spatial-resolution HRTF measurements for 45 different subjects” [1]

