## Supplementary Figures for "Two Are Better Than One: Solving the Problem of Vertical Sound Source Localization via Binaural Integration of HRTFs"

### Supporting Information

#### S1 Figure

##### Correlation of learned map with HRTFs.

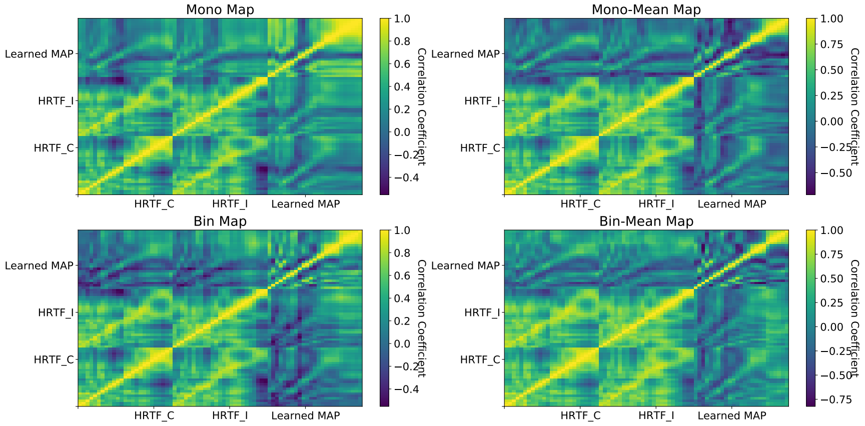


**Figure 8. Similarity of HRTFs.** Cross-correlation of HRTFs with learned map for single participant. Comparison of the learned map under different conditions with the HRTFs of the left and right ear, respectively, via a cross-correlation. Top left shows comparison to a map learned using monaural inputs. Top right depicts comparison to a map learned using monaural inputs with integrated prior information. Bottom left shows comparison to a binaurally learned map. Bottom right depicts comparison to a map learned using binaural inputs with integrated prior information. Colors indicate the correlation coefficient index.

#### S2 Figure

**Localization performance for different signal-to-noise ratios.**

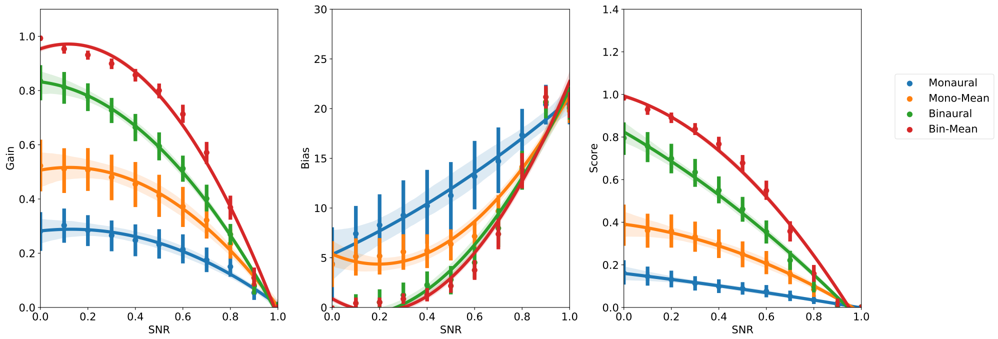


**Figure 9. Signal to noise ratio test.** Influence of signal-to-noise ratio on localization performance is depicted. The learned map is based on binaural signals integrating prior information. Left panel, the averaged gain values for the linear regression over all participants is shown. Middle panel, the bias values for the linear regression over all participants is depicted. Right panel, coefficient of determination for the linear regression over all participants is shown. The bars indicate the confidence interval (95%). Different line colors indicate various tested conditions as described in Results. Blue lines represent localization of monaural sings. Orange lines show localization for monaural signals integrating prior information. Green lines indicate localization for binaural signals. Red lines represent localization for binaural signals integrating prior information.

#### S3 Figure

**Setup for behavioral experiment.**

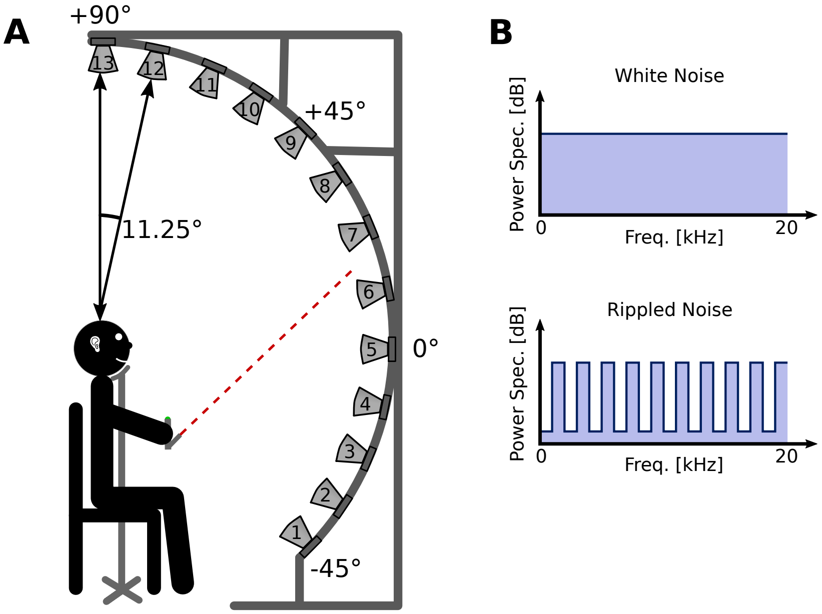


**Figure 10. Behavioral Setup. A** Experimental setup used for the behavioral experiment. Participants' head is placed on a chin rest with equal distance to the speakers (120cm). Participants used a laser pointing device to indicate from where they perceived a sound. **B**
Frequency representation of white noise stimulus (upper plot). Frequency representation of rippled noise stimulus (lower plot).
